# Advancing the genetic utility of pre-clinical species through a high-quality assembly of the cynomolgus monkey (*Macaca fascicularis*) genome

**DOI:** 10.1101/2020.05.01.072280

**Authors:** Elias Oziolor, Shawn Sullivan, Hayley Mangelson, Stephen M. Eacker, Michael Agostino, Laurence Whiteley, Jon Cook, Petra Koza-Taylor

## Abstract

The cynomolgus macaque is a non-human primate model, heavily used in biomedical research, but with outdated genomic resources. Here we have used the latest long-read sequencing technologies in order to assemble a fully phased, chromosome-level assembly for the cynomolgus macaque. We have built a hybrid assembly with PacBio, 10x Genomics, and HiC technologies, resulting in a diploid assembly that spans a length of 5.1 Gb with a total of 16,741 contigs (N50 of 0.86Mb) contained in 370 scaffolds (N50 of 138 Mb) positioned on 42 chromosomes (21 homologous pairs). This assembly is highly homologous to former assemblies and identifies novel inversions and provides higher confidence in the genetic architecture of the cynomolgus macaque genome. A demographic estimation is also able to capture the recent genetic bottleneck in the Mauritius population, from which the sequenced individual originates. We offer this resource as an enablement for genetic tools to be built around this important model for biomedical research.

## Introduction

The origins and genetic diversity in cynomolgus macaques (*Macaca fascicularis*) are complex and heavily examined questions. Cynomolgus macaques are distributed across mainland Asia [1], as well as isolated island populations, such as Mauritius [2] and the Philippines [3]. While the geographic origin of cynomolgus macaques is largely contested with a variety of evidence from genetic studies [3-6], there is clear support showing that isolated island populations stem from the Indochinese region [7]. The Mauritius cynomolgus macaque populations are genetically isolated from other island populations [8] and exhibit substructure even within the island [8-10]. The macaques on the Mauritius are largely bottlenecked with an estimated ∼20 founding individuals [11].

Despite their genetic isolation and largely limited population diversity, Mauritius cynomolgus macaques have become an important source for non-human primate studies in biomedical research [12]. Macaque physiology is both well documented and highly homologous to that of human [13]. Consequently, cynomolgus macaques are often used as disease models [13]. Cynomolgus and rhesus macaques *(Macaca mulatta)* have a long history of ancestral introgression [14, 15]. While rhesus macaques also have comparable homology with humans in both physiology and genetics [15, 16] cynomolgus macaques have surpassed rhesus in the number of imported animals for biomedical research [8, 12].

Sequencing of the cynomolgus macaque genome has allowed for an unprecedented ability to perform genetic comparisons [16]. It has enabled studies of ancestry [6] and has added to the utility of this species as a pre-clinical model [15-17]. Few examples exist of the utility of these data in drug safety [18-20]. There have been several assemblies of the cynomolgus macaque genome stemming from individuals from Malaysia [21], Vietnam [16] and Mauritius [22]. While highly novel at their time, these assemblies were largely made with the available short-read sequencing technology, popular in the early 2010s. The genomes’ contiguity and accuracy are heavily impacted by the lack of long-read sequences, resulting in limited utility.

Long-read sequencing technologies are rapidly emerging and providing opportunities for exciting sequencing studies. Tools for analyzing long-read data are constantly emerging [23] and many are becoming widely accepted in medical genetics [24], as well as key to discovering novel clinically important variants [25]. However, one of the most essential functions of long-read technology was immediately recognized in their ability to vastly improve the contiguity of genome assemblies [26]. Long-read genomes that have immerged for various non-human primates, such as gorillas (*Gorilla gorilla*) [27], rhesus macaques [28], chimps (*Pan troglodytes*) and orangutans (*Pongo abelii*) [29], as well as golden snub-nosed monkeys (*Rhinopithecus roxellana*) [30], showcase the incredible advancements in the quality of genomes that can be made with the advent of modern sequencing.

Given the wide use of cynomolgus macaques in biomedical research, as well as their relatively sub-par genome assemblies compared to other primates, we argue that it is imperative to create a higher-quality assembly. In this manuscript, we used a combination of PacBio, 10x Genomics, HiC and Illumina HiSeq technologies to create a high-quality, phased assembly of the cynomolgus macaque genome. We created a hybrid assembly and compared it to related species, as well as to the previous versions of the cynomolgus macaque genome. Here, we provide a crucial advance in the utility of this important pre-clinical species as a genetic resource for advancement of biomedical research.

## Methods/Results

### Sample collection and sequencing

The cynomolgus macaque used for this study was farm bred by Bioculture (Mauritius) Ltd., born on Feb 26, 2012 and arrived at Charles River Lab on Nov 17^th^, 2015. It was housed by Comparative Medicine in Groton, CT from May 20^th^, 2016 to Oct. 21, 2019, where it underwent acclimation, further screening and maintained SPF status.

We collected blood into a K2 EDTA vacutainer, inverted a few times and kept at 4°C. We extracted high molecular weight DNA using magAttract HMWDNA Kit (Qiagen 67563). We assessed quantity by Nanodrop (Thermo Scientific) and quality by low agarose gel electrophoresis and pulse field gel electrophoresis (Novogene). The DNA was > 50Kbp, stored at 4°C and shipped in dry ice to Novogene Inc. for sequencing. Library preparation proceeded using standard methods for PacBio, 10x and Illumina HiSeq. Sample was sequenced at high depth to ensure sufficient coverage and error correction (Table 1).

**Table 1:** Sequencing technologies and depth used for hybrid cynomolgus macaque assembly.

| Library type | Insert size | Total data (Gb) | Read length (bp) | Sequence coverage(X) |
| --- | --- | --- | --- | --- |
| Illumina HiSeq | 350 | 315.37 | 150 | 104.8 |
| PacBio |  | 92.92 |  | 30.67 |
| 10X Genomics | 600 | 223.94 | 150 | 73.91 |
| HiC |  | 163.40 | 150 |  |

Blood was also sent to Phase Genomics (Seattle, WA) where chromatin conformation capture data was generated using a Phase Genomics Proximo Hi-C 2.0 Kit, which is a commercially available version of the Hi-C protocol [31]. Intact cells were crosslinked using a formaldehyde solution, digested using the Sau3AI restriction enzyme, ends filled in with biotinylated nucleotide, and proximity-ligated to create chimeric molecules composed of DNA fragments that were physically proximal *in vivo* but not necessarily genomically proximal. These chimeric molecules were then pulled down by streptavidin beads and used to generate an Illumina-compatible sequencing library. An Illumina HiSeq4000 was used to generate a total of 544,671,413 PE150 read pairs.

### De novo *assembly*

The FALCON de novo assembly, FALCON-Unzip haplotype segregation, and FreeBayes short read polishing with FreeBayes were performed with DNAnexus internal workflows on the DNAnexus platform. The FALCON (falcon-2018.31.08-03.06-py2.7-ucs4-beta release) and FALCON-Unzip binaries (falcon-2018.04.06-08.39-py2.7-ucs4-alpha release) were obtained from the Pacific Biosciences cloud server (https://downloads.pacbcloud.com/public/falcon/).

First, single-molecule, real-time sequencing (SMRT) data at approximately 30.67x coverage was passed through the TANmask and REPmask modules from the Damasker suite [32]. These tools mask repetitive regions for use during the initial read overlap discovery, causing the overlapper to instead focus on unique sequences to identify reads that overlap because they originate from the same genomic locus. This masked data was then used as input to the Daligner stage of the traditional FALCON pipeline, using a read-length cutoff of 2000bp to error-correct the raw-reads. Next, the initial contigs were again passed through the TANmask and REPmask modules, followed by the Daligner overlap portion of the FALCON pipeline. For this second overlap portion of the FALCON pipeline, a length cut-off of 1 bp was selected. Finally, these contigs were passed to the FALCON assembly stage of the DNAnexus workflow, which converts these contigs to an assembly graph, assembles the contigs using the graph, then identifies alternative contigs. The resulting *de novo* assembly was separated into haplotype contigs (haplotigs) using the FALCON-Unzip workflow on the DNAnexus platform, leveraging a polishing step from PacBio’s Arrow algorithm (SMRT Link v6.0.47841). Outputs from FALCON-Unzip consisted of primary contigs and haplotigs. These were concatenated, then Illumina reads (104.8x coverage) were aligned to the combined assembly using BWA MEM [33] v0.7.15-r1140. The short-read polishing was then performed using FreeBayes v1.2.0-17-ga78ffc0 with the BAM file from BWA and assembly contigs [34].

Hi-C reads were aligned to the polished FALCON-Unzip assembly using BWA-MEM [33] with the −5SP and -t 8 options specified, and all other options used default settings. SAMBLASTER [35] was used to flag PCR duplicates, which were later excluded from analysis. Alignments were then filtered with samtools [36] using the -F 2304 filtering flag to remove non-primary and secondary alignments.

FALCON-Phase [37] was used to correct likely phase switching errors in the primary contigs and alternate haplotigs from FALCON-Unzip and output its results in pseudohap format, creating one complete sets of contigs for each phase.

Phase Genomics’ Proximo Hi-C genome scaffolding platform was used to create chromosome-scale scaffolds from the FALCON-Phase phase 0 assembly, following the same single-phase scaffolding procedure described in Bickhart et al. [26]. As in the LACHESIS method [38], this process computes a contact frequency matrix from the aligned Hi-C read pairs, normalized by the number of Sau3AI restriction sites (GATC) on each contig and constructs scaffolds that optimize expected contact frequency and other statistical patterns in Hi-C data. Approximately 20,000 separate Proximo runs were performed to optimize the number of scaffolds and scaffold construction in order to make the scaffolds as concordant with the observed Hi-C data as possible. This process resulted in a set of 22 chromosome-scale scaffolds containing 2.41 Gb of sequence (99.97% of the corrected assembly).

Juicebox [39] was then used to correct scaffolding errors, and FALCON-Phase [37] was run a second time to detect and correct phase switching errors that were not detectable at the contig level, but which were detectable at the chromosome-scaffold level. Metadata generated by FALCON-Phase about scaffold phasing was used to generate a Juicebox-compatible assembly file for each phase indicating identical order and orientation of contigs within scaffolds. These files were subsequently used to produce a diploid, fully phased, chromosome-scale set of scaffolds using a purpose-built script (https://github.com/phasegenomics/juicebox). Gap closing was performed on each phase by PBJelly2 from PBSuite v15.8.24 [40] using the same 30.67x PacBio long reads as were used in the initial FALCON assembly. Benchmarking Universal Single-Copy Orthologs (BUSCO) v3.0.2 [41] was used to assess genome completeness using the Eukaryota obd 9 dataset.

Repetitive elements were identified and classified by RepeatModeler v1.0.11 [42] and RepeatMasker v4.0.9 [43] relative to Repbase-derived RepeatMasker libraries v20181026 [44]. The MAKER v2.31.10 [45] pipeline was used to perform whole-genome annotation with expressed sequence tag (EST) evidence in the form of cDNA sequences from Macaca fascicularis (*Macaca_fascicularis*_5.0) available through the Ensembl database [46], Ensembl cDNA sequences from Homo sapiens (GRCh38.p13) and aligned ESTs from *Macaca fascicularis* [17, 30]. Protein homology evidence was provided by the uniprot_sprot database (downloaded 11/6/2019), and Ensembl protein sequences for *Homo sapiens* (GRCh38.p13) and *Macaca fascicularis* (*Macaca_fascicularis*_5.0). The consensi.fa.classified output from RepeatModeler was used for soft repeat-masking of the assembly. Finally, MAKER was provided an Augustus gene prediction model for *Macaca fascicularis* developed by BUSCO v3.0.2 [41]. After MAKER output was generated, the uniprot_sprot database was used to give functional names to annotations.

### Genome contiguity and accuracy

The genome produced is a fully phased, chromosome scale assembly of the cynomolgus monkey genome. The diploid assembly spans a length of 5.1 Gb with a total of 16,741 contigs (N50 of 0.86Mb) contained in 370 scaffolds (N50 of 138 Mb) positioned on 42 chromosomes (21 homologous pairs). There are 16371 gaps that account for 0.0093% (0.47 Mb) of the entire of the genome. The two phases are similar in length, though Phase 0 is 0.0036% (92.8 Kb) larger than Phase 1. Phase 1 also contains one more contigs (8371) than Phase 0 and, subsequently, one additional gap (8186). The contig N50 for each phase is 0.86 Mb and the scaffold N50 is 138 Mb, though the Phase 0 scaffold N50 is 0.018% (24.5 Kb) larger. Gene space completeness was assessed by BUSCO, which uses a core of highly conserved orthologous genes (COGs) to quantify functional gene content. Of the 303 BUSCO groups searched, 212 (70.0%) were identified as complete with another 24 (7.9%) classified as fragmented (Complete: 70.0% [Single-Copy: 65.0%, Duplicated: 5.0%], Fragmented: 7.9%, Missing: 22.1%).

We aligned Phase 0 to the most current genome assembly from RefSeq (cynomolgus_macaque_v5) using MUMmer4 [47] (Figure 2). Of Phase 0, 98.9% of sequences mapped successfully to cynomolgus_macaque_v5 and 99.2% of bases successfully mapped. Over 9 million SNPs were identified between the two genomes, which was unsurprising given the sequences were derived from two non-human primates. Of all sequences, 2.46 Gb aligned one to one between the two genomes, suggesting high sequencing concurrence (Figure 2). The new assembly successfully placed more sequences in chromosomal designations and on fewer scaffolds (185 vs. 7601), suggesting substantially improved contiguity (Figure S1).

**Figure 1:**
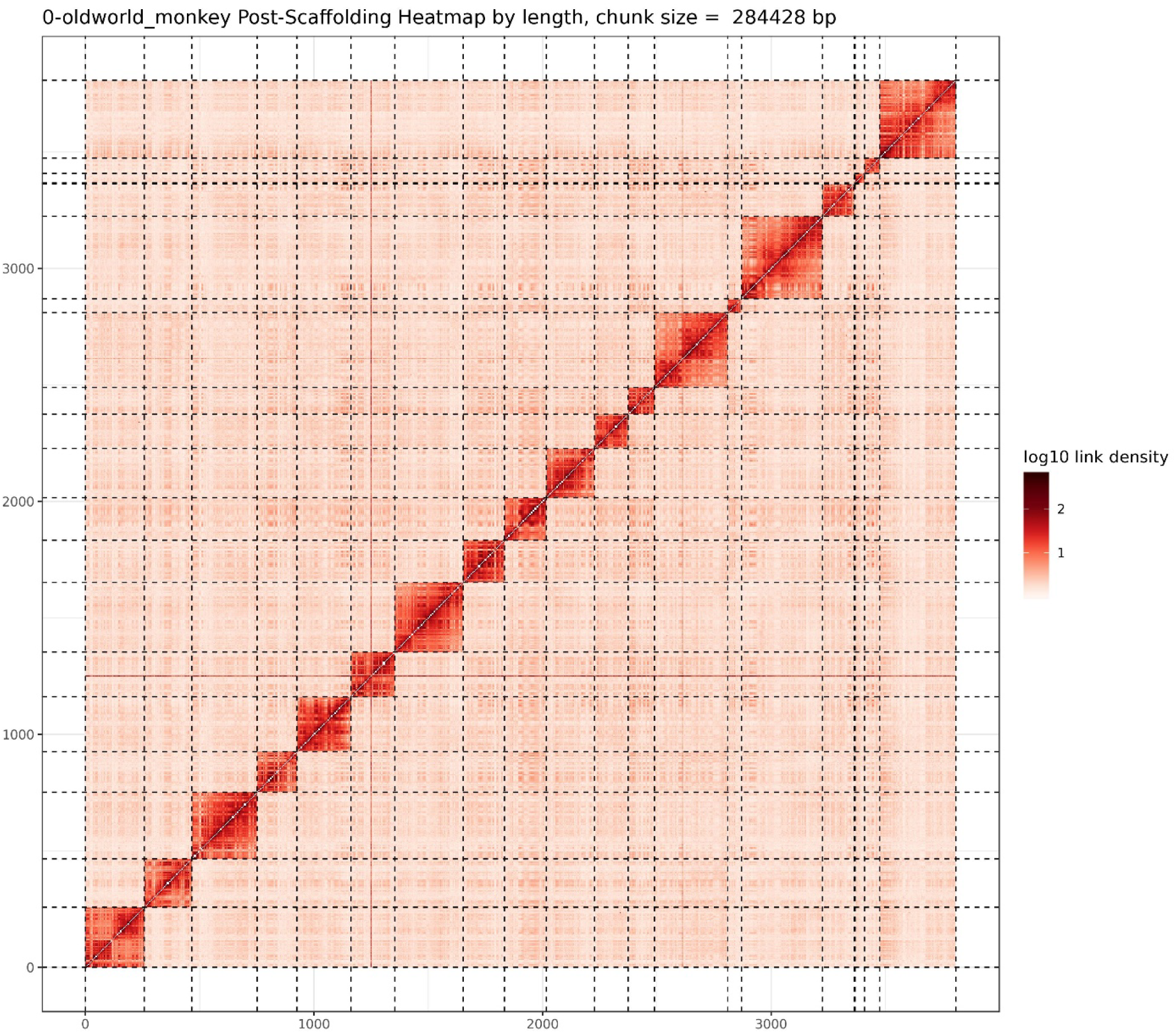
HiC post-scaffolding heat map of linkage groups reveals high link density in groups suggesting strong chromosomal designations.

**Figure 2:**
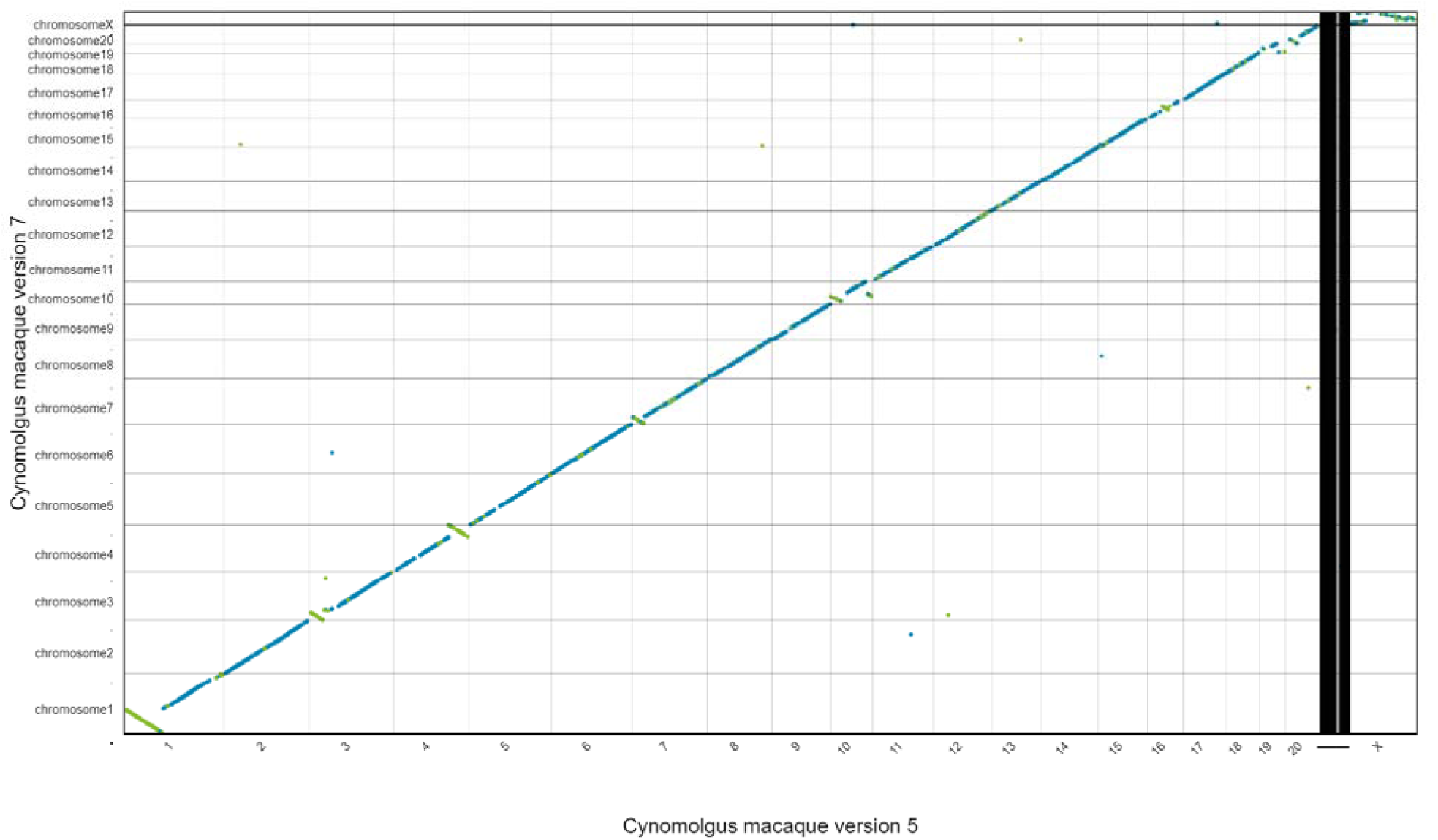
Alignment to previous public version of the cynomolgus macaque suggests strong contiguity, with multiple inversions identified across original chromosomes.

To visualize the assembly, we performed several comparisons of our scaffolds with various other assemblies using D-GENIES [48]. First, we compared our Phase 0 scaffolds to our Phase 1 scaffolds and observed clear linear correspondence between the two phases, suggesting no significant structural variation between the two phases (Figure S3). We then compared the Phase 0 scaffolds to the human reference genome hg38 [49] (Figure S4). Most monkey scaffolds in Phase 0 had a clear homologous human chromosome mate, although many structural variants are visible. Two exceptions were the monkey chromosomal scaffold named Contig4, which contains regions mapping to human chromosomes 7 and 21, and monkey chromosomal scaffolds Contig2 and Contig16, which both contain regions mapping to human chromosome 2. We observe high contiguity between the Phase 1 scaffolds to *M. mulatta* genome (Figure 4). We observe only two major re-assignments of >80kb scaffolds to different chromosomes. However, we detected high contiguity in regions where our assembly differs structurally from *M. fascicularis* (ex. Chromosomes 3, 5, 10), suggesting we may have resolved a formerly mis-identified structural variant (Figure S5). Other inversions coincide with inversions between our assembly and the former *M. fascicularis* (Chromosomes 1, 4, and 7). Given the high spatial resolution of our assembly, we have high confidence in the structural variability we have identified.

**Figure 3:**
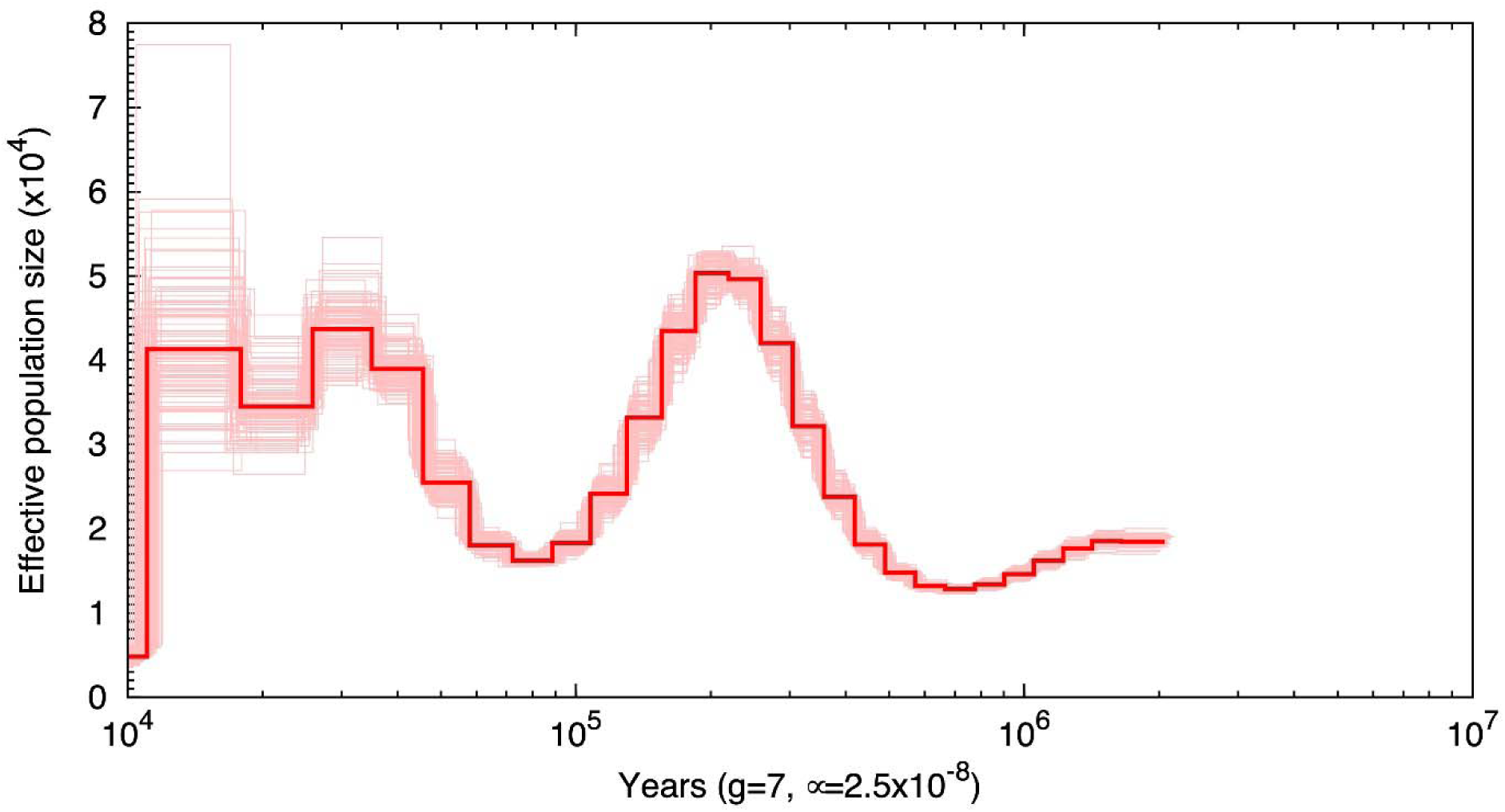
Effective population size (2N_o_) swings widely with time, likely due to population expansion in the deep history of cynomolgus macaques. The most recent period of low population size estimates reflects the known bottleneck of cynomolgus macaques on the Mauritius island.

**Figure 4:**
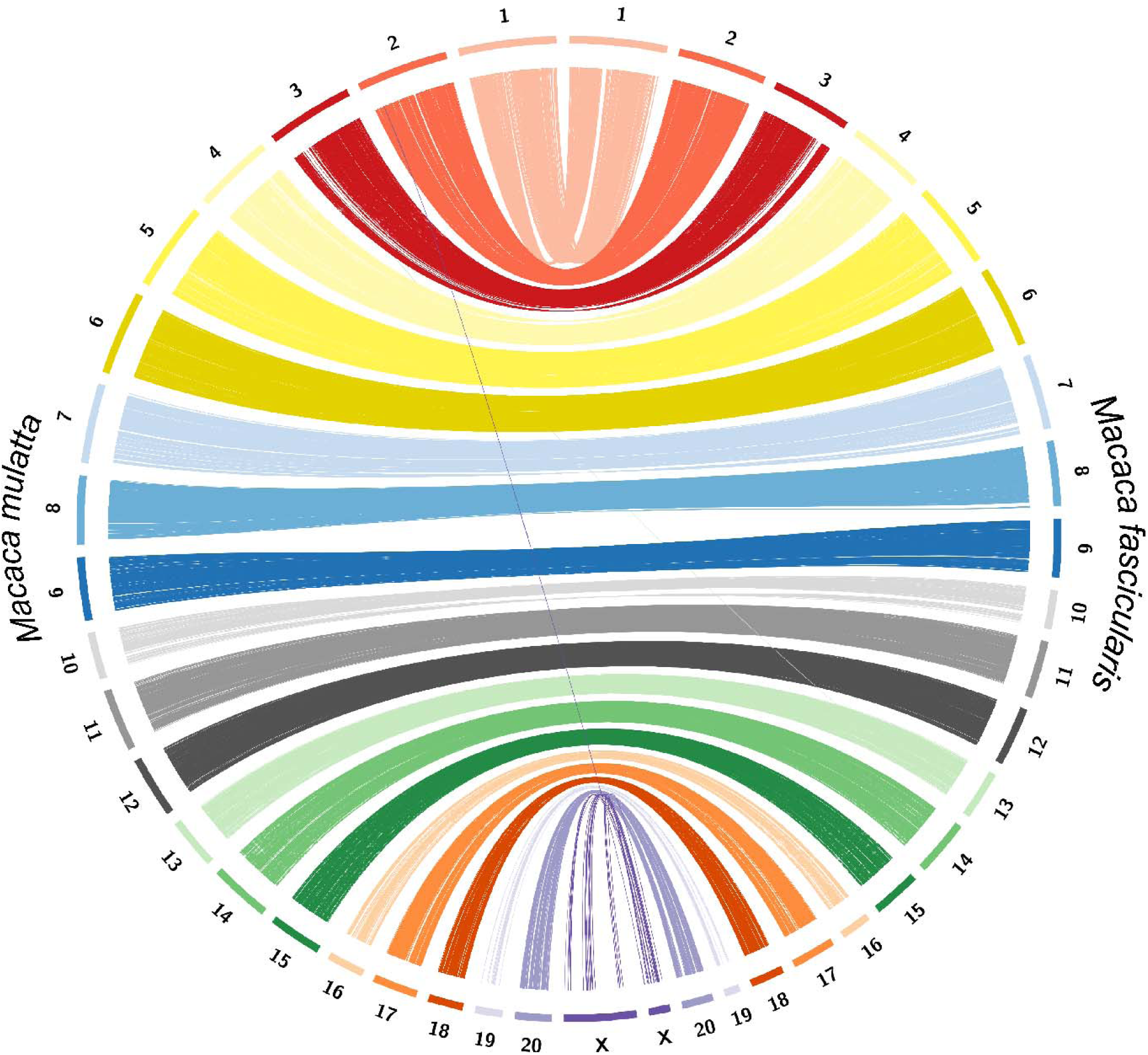
Circos plot of genomic synteny regions across *M. fascicularis* and *M. mulatta*. The high congruence of genome assemblies supports the close phylogenetic relationship between the two species and suggests tight linkage on larger genomic regions. Only two genomic re-assignments are observed with size above 80 Kb. One of them maps to the more poorly assembled X chromosome, while the other is a re-assignment of a region on from Chromosome 3 to 13 in *M. fascicularis*.

An additional novel assembly(v6) of the cynomolgus macaque genome has become publicly available (RefSeq: GCA_011100615.1). We repeated the MUMmer4 alignment of our Phase 0 assembly to the cynomolgus_macaque_v6. We show that our hybrid assembly identifies structural variations in common with both v5 and v6 assemblies that are not captured in previous assemblies but are likely correct because of the advantage of HiC sequencing in parsing complex structural changes (Figure S2). Using methods originally shown in LACHESIS [38], we can compute the log-likelihood of a contig having generated the observed Hi-C data given its orientation and neighboring contigs. This method provides an internal heuristic measurement for assessing the quality of a contig’s placement in the scaffolds for evaluating *de novo* scaffolds without external data sources. Proximo considers a contig to be in a high-quality orientation when the log-likelihood of the observed Hi-C data having been generated is at least 100. In both the Phase 0 and Phase 1 scaffolds, 93.3% of the scaffolded length was scaffolded in a high-quality orientation, with the remainder being in a positive log-likelihood orientation but below the threshold of 100.

### Assembly accuracy

To assess the accuracy of our assembly, we took the Illumina HiSeq paired end reads (∼315Gb) and mapped them to the newly assembled genome. We used trimmomatic version 0.39 [50] to retain appropriate read pairs with high quality phred scores (see GitHub for specific code used). We mapped reads onto the new assembly using Burrows-Wheeler Aligner version 0.7.17 with the MEM algorithm [33] and filtered on mapping quality with samtools version 1.9 [36]. We merged alignments from multiple lanes using bamtools version 2.5.1 [51] and called variants using freebayes version 1.3.1-17 [34]. Of the short reads 99.08% mapped with proper pairs to the genome and using bedtools [52], we estimated that the HiSeq reads covered 95.9% of the assembly.

### Demographic history

We estimated the historic demographic history of cynomolgus macaques from the Mauritius individual genome sequence using a Pairwise Sequentially Markovian Coalescence (PSMC) approach [53]. We used samtools mpileup followed by bcftools version 1.10.2 [54] to create a consensus sequence for our individual using aligned Illumina HiSeq reads. We estimated population demography with default parameters, with exception to generation time, for which we used an estimate of 7 years as per estimates from other macaque studies [21] and iterated the algorithm 100 times with replacement to explore variation in estimated size. The population size of cynomolgus macaques had upwards and downwards swings over time, likely due to expansion or hybridization events, followed by bottlenecks (Figure 3). A most prominent expansion event occurred ∼20,000 years ago, almost quadrupling the effective population size of cynomolgus macaques. A much more recent and rapid bottleneck (∼1000 years ago) event is detected by PSMC, which is likely reflective of the geographic and genetic isolation of cynomolgus macaques on the Mauritius islands. The estimate shows that the effective population size of cynomolgus macaques has dropped from ∼4500 to ∼480 drastically, thus retaining only 10% of its original variation (estimated theta=0.17). These data showcase a severe genetic isolation, which is apparent even in methods designed to examine deep demographic history.

## Conclusions

The cynomolgus macaque is a keystone species for biomedical research. The main draw for cynomolgus macaques is their well-documented physiology. The molecular resources for continuing to use cynomolgus macaques in research are highly dependent of the quality of the cynomolgus genome available. Here we present a high-quality, phased hybrid genomic assembly with chromosome length scaffolds as a result of deep sequencing with short, long and linked read technologies. Our assembly correctly identifies structural variations not present in current cynomolgus macaque assemblies. We offer this assembly as a public resource to enable high level molecular and genetic explorations of the cynomolgus macaque as a continued model for biomedical research.

## Data and code

The code used for analyses in this manuscript can be found at the following GitHub repository: https://github.com/eoziolor/mfascicularis_v7. The assembly and annotation can be found in the European Nucleotide Archive (Sequencing accession numbers: GCA_903645265, GCA_903645275).

## Suplementary

**Figure S1:**
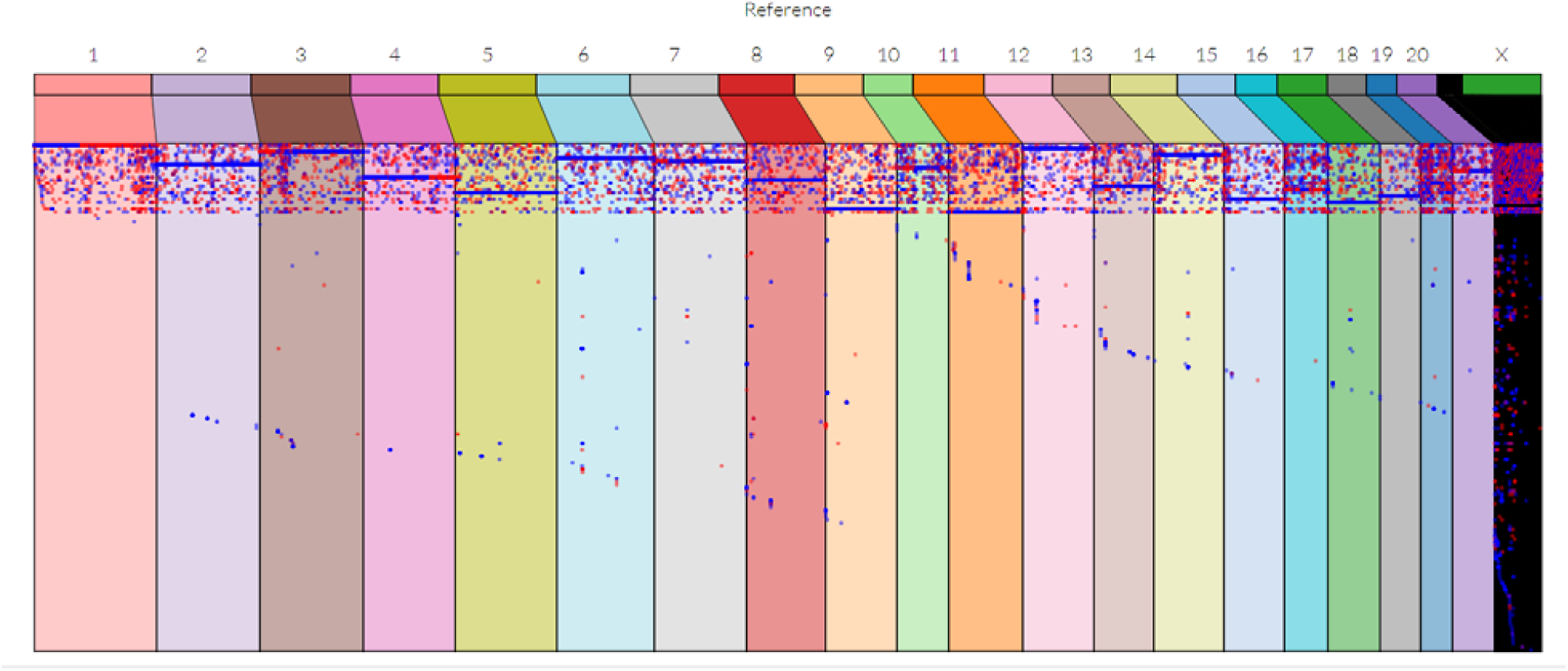
Shift in chromosome windows reveals higher number of reads placed on chromosomal designations within this new assembly.

**Figure S2:**
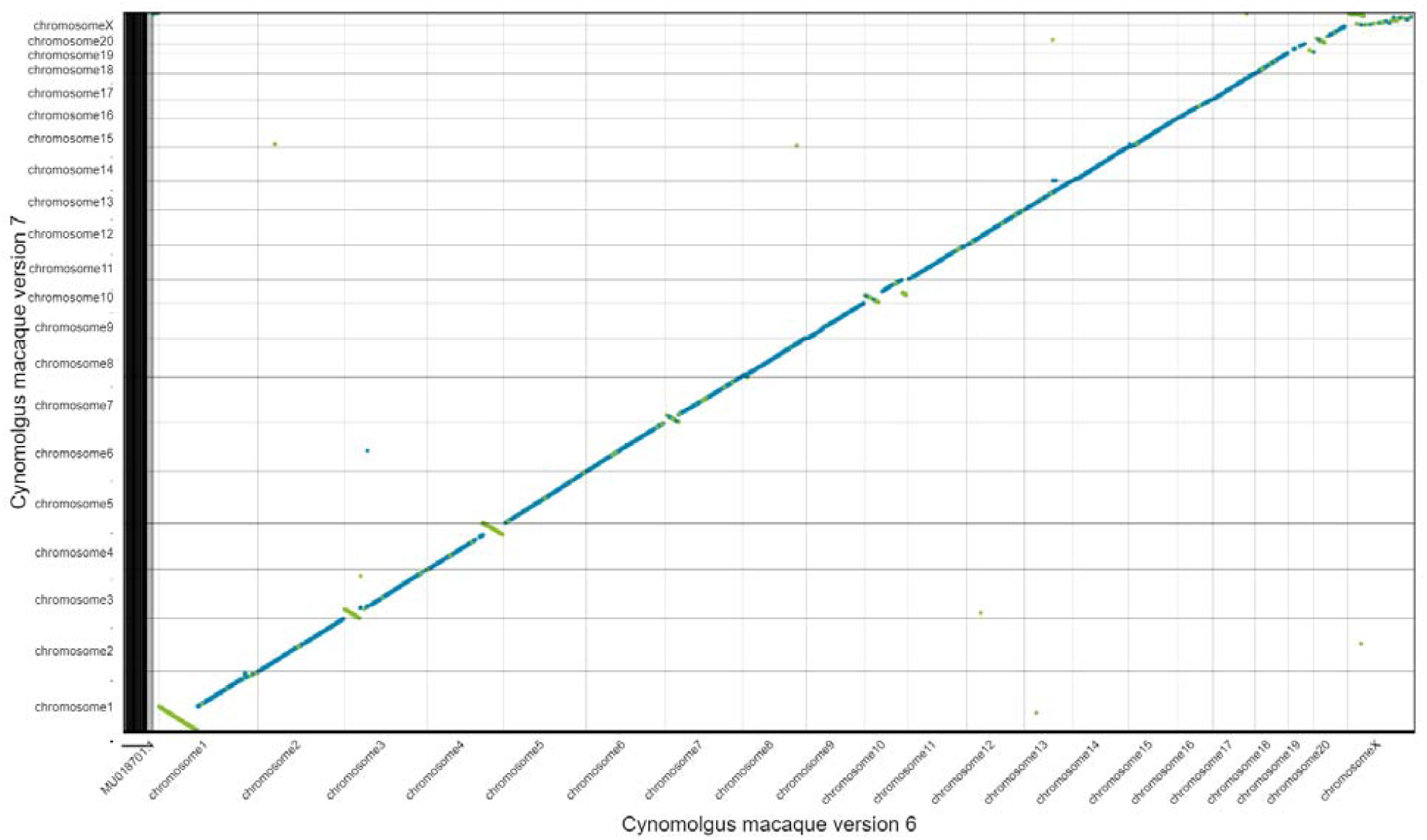
Alignment of cynomolgus macaque assembly, version 6, with our new hybrid assembly reveals structural variations that are novel to our assembly, but commonly mis-identified in both previous assemblies.

**Figure S3:**
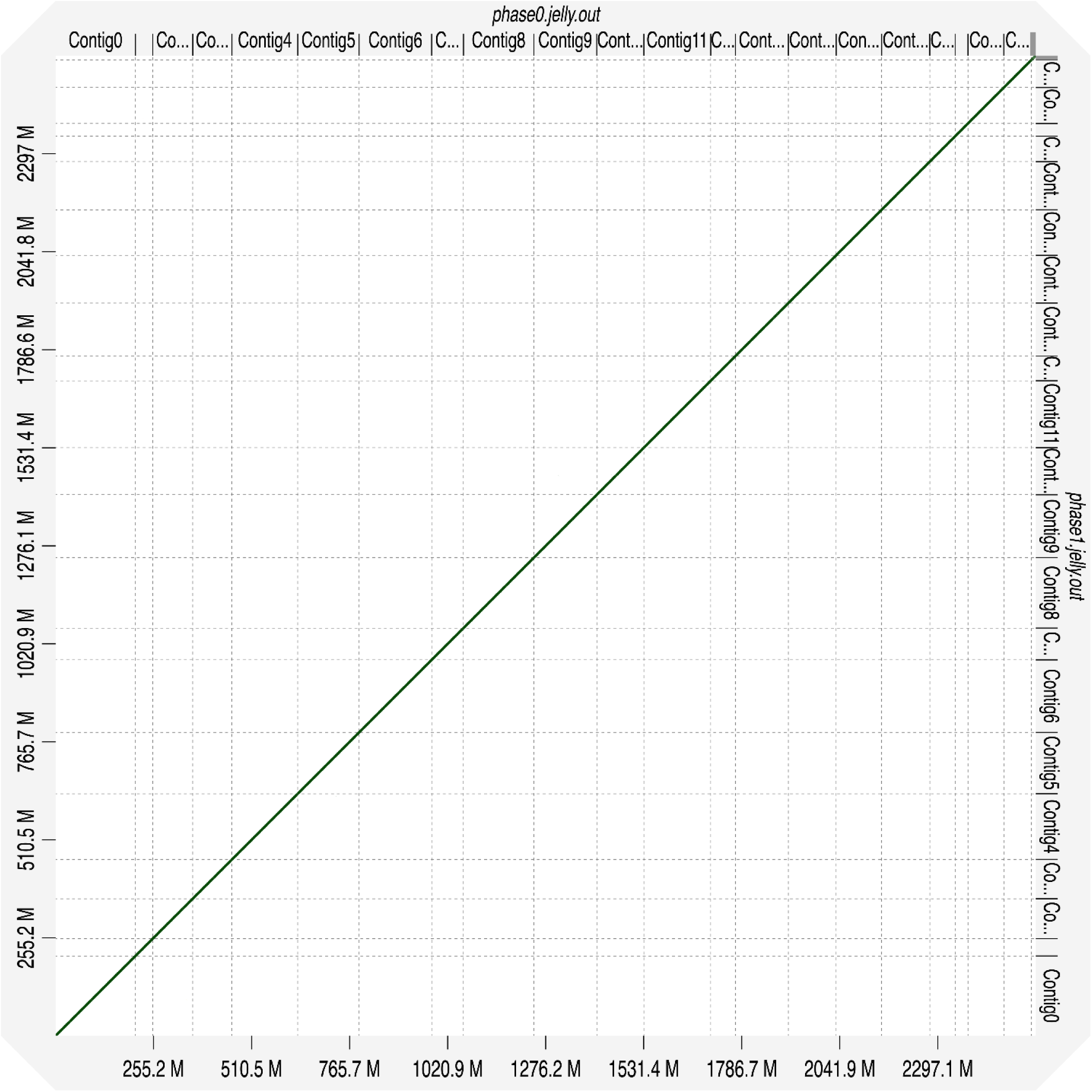
Alignment of cynomolgus monkey Phase 0 scaffolds to Phase 1 scaffolds shows no significant structural variation between phases.

**Figure S4:**
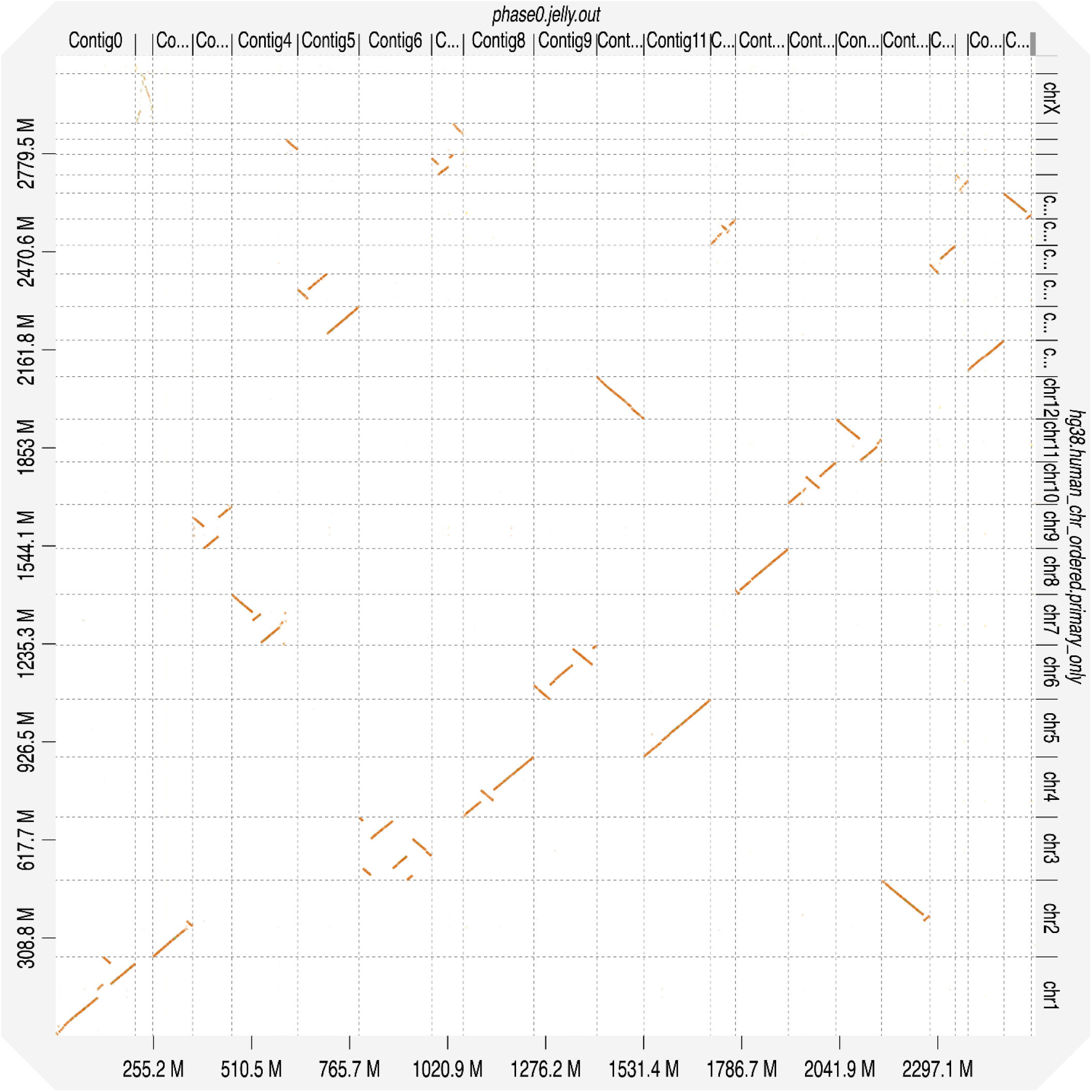
Alignment of cynomolgus monkey Phase 0 scaffolds to the human reference genome hg38, with the “Strong Precision” option set in D-GENIES. Numerous structural variations are present, though most monkey chromosomes have a clear homologous matching human chromosome. Of note, the monkey chromosome scaffold named Contig4 contains regions mapping to two human chromosomes, chromosomes 7 and 21, and human chromosome 2 contains regions mapping to two monkey chromosomes, Contig2 and Contig16.

**Figure S5:**
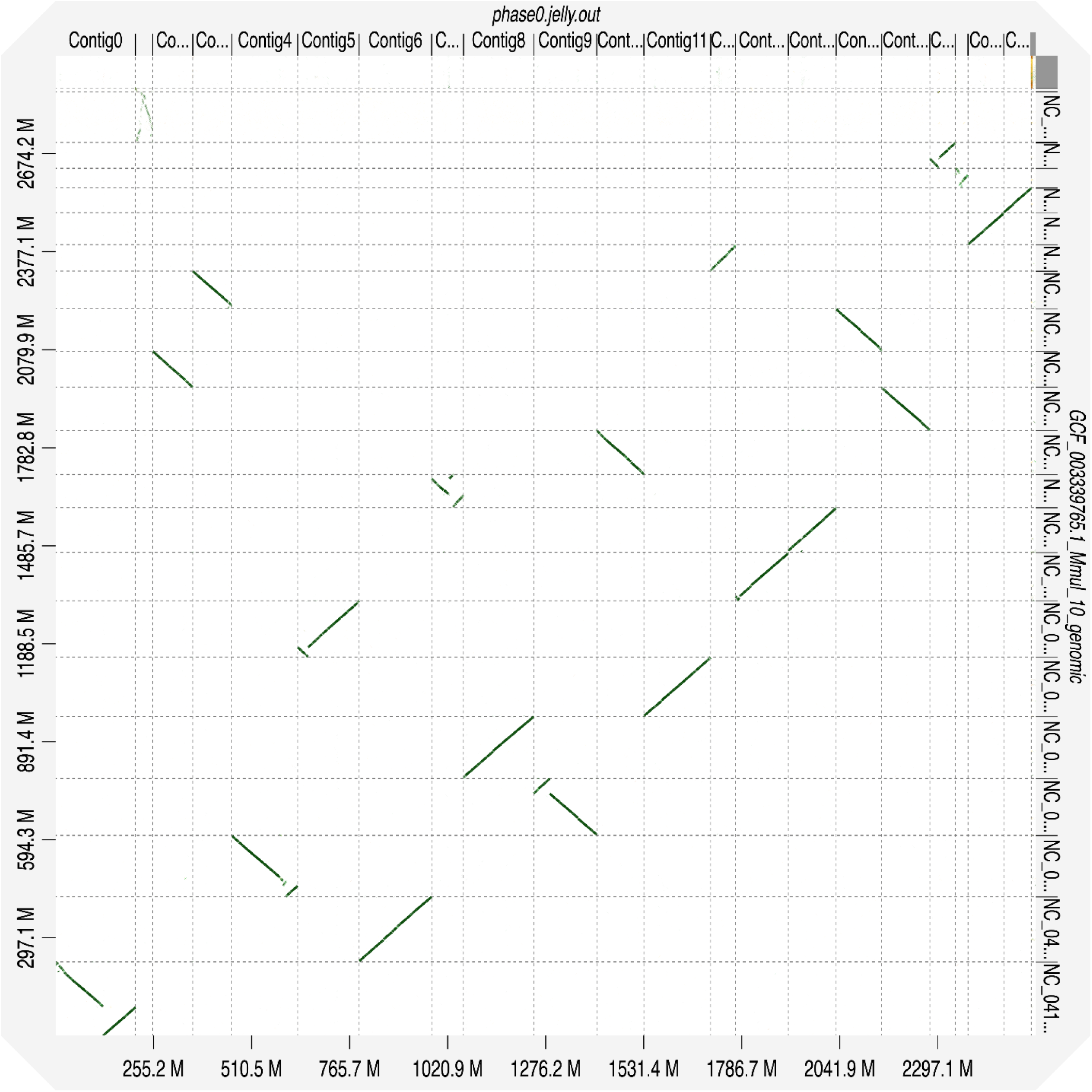
Alignment of cynomolgus monkey Phase 1 scaffolds to the reference genome of *M. mulatta*, with the “Strong Precision” option set in D-GENIES. Large portion of structural variant patterns observed comparing the Phase 0 scaffolds to cynomolgus macaque seem to be observed here as well. This may lead to better understanding of previously poorly structured regions in both monkey genomes.

